# Docosahexaenoic acid alone and in combination with carboplatin significantly reduces tumor cell growth in preclinical models of ovarian cancer

**DOI:** 10.1101/2020.11.10.376152

**Authors:** Olena Bilyk, Bahareh Hamedi, Indrani Dutta, Marnie Newell, Amirali B. Bukhari, Armin M. Gamper, Rojine C. McVea, Jiahui Liu, Julia Schueler, Gabrielle M. Siegers, Catherine J. Field, Lynne-Marie Postovit

## Abstract

Despite recent advances in diagnosis and treatment, ovarian cancer (OC) is the most lethal gynecological malignancy and improving the efficacy of chemotherapy is of great interest. This study increases our understanding of how dietary intervention with docosahexaenoic acid (DHA)-a supplement proven safe for human consumption – enhances the anti-cancer effects of conventional chemotherapy. Our results demonstrated synergistic cell killing by DHA and carboplatin in OC cell lines. Furthermore, DHA supplementation alone and in combination with carboplatin significantly reduced OC growth in a high-grade serous OC patient-derived xenograft mouse model. Carboplatin administered intraperitoneally significantly reduced tumor growth in DHA-fed mice compared to mice on the control diet. Intravenous carboplatin administration in combination with DHA reduced tumor growth similarly to carboplatin or DHA monotherapies. The DHA-induced reduction in tumor growth in this model was associated with increased tumor necrosis and improved survival. As such, our findings provide a strong rationale to move to clinical trials that will determine whether DHA supplementation enhances the efficacy and tolerance of cytotoxic chemotherapy in patients with OC.

## INTRODUCTION

Ovarian cancer (OC) is the most lethal gynecological malignancy in the western world. Despite recent advances in diagnosis and treatment, the 5-year survival rate is still poor, especially in patients with advanced disease. This is particularly true for epithelial OC, which accounts for approximately 90% of all OC diagnoses (Orr & Edwards, 2018; Mallen *et al,* 2018). Due to subtle symptoms of the early-stage disease and a lack of effective screening methods, most patients present with advanced stages. Cytoreductive surgery, in combination with adjuvant or neoadjuvant platinum/taxol-based chemotherapy, is the first line treatment approach in most cases. While their initial response rate is typically good, the majority of patients relapse with chemoresistant disease, which results in treatment failure and poor overall survival (Bristow *et al*, 2009; Pignata *et al*, 2017; Pokhriyal *et al*, 2019). As such, further advances in research are clearly needed to improve treatments and extend the lives of women affected by OC.

Omega-3 long chain polyunsaturated fatty acids (N-3 LCPUFA), including docosahexaenoic acid (DHA), are widely recognized as dietary components that can improve human health (Sambrook *et al*, 2009). Previous results have shown that beneficial health effects of N-3 LCPUFA, particularly DHA, are mediated through alterations in the composition of fatty acids in membrane-associated lipid rafts. By altering lipid raft composition, DHA can facilitate the association of myriad signaling molecules with receptors, ultimately driving the expression of several genes involved in cell survival and death (Ma *et al*, 2004; Blanckaert *et al*, 2010; Corsetto *et al*, 2011).

There is a strong rationale for investigating nutritional interventions, such as N-3 LCPUFA, in cancer prevention and therapy (Sawyer & Field, 2010). In OC, although the therapeutic effects of N-3 LCPUFA have not been extensively studied, several *in vitro* and *in vivo* reports have shown that DHA can induce apoptosis and cell cycle arrest (Han *et al*, 2016), reduce cell growth (Tanaka *et al*, 2017; Sharma *et al*, 2005), invasion and metastasis (Wang *et al*, 2016), and induce sensitivity to cisplatin *via* multiple molecular pathways (Zajdel *et al*, 2018). Preclinical studies have shown an association between incorporation of DHA into tumor membrane phospholipids and increased efficacy of chemotherapy against different types of cancer. Examples include enhanced effectiveness of doxorubicin in lung cancer (Hardman *et al*, 2000), mitomycin-C in colorectal cancer (Yang *et al*, 2013), doxorubicin, vincristine and fludarabine in chronic lymphoid leukemia (Fahrmann & Hardman, 2013), 5-FU in gastric cancer (Gao *et al*, 2016) and doxorubicin, herceptin and docetaxel in breast cancer (Ewaschuk *at el*, 2012; Newell *et al*, 2019). Clinical studies have reported that dietary intake of DHA results in increased DHA incorporation in breast adipose tissue that correlates with enhanced responses to chemotherapy and improved patient outcomes (Yee *et al*, 2010; Bougnoux *et al*, 2009). Moreover, supplementation with DHA at a wide range of doses reduces the occurrence and severity of adverse events caused by chemotherapeutic drugs in a range of malignancies including breast, lung, pancreatic and colorectal cancers (Bougnoux *et al*, 2009; Morland *et al*, 2016).

Herein, we have used multiple cell lines to establish the efficacy of DHA, alone and in combination with chemotherapy, in reducing the growth of OC *in vitro*. Furthermore, we have extended these studies *in vivo*, using a patient derived xenograft (PDX) model that recapitulates human disease (Go & Yoshida, 2020).

## RESULTS

### DHA has differential effects on cell viability and apoptosis in ovarian cancer cell lines and normal immortalized cells

Using MTT and crystal violet assays, we assessed metabolic activity and cell viability of four OC cell lines after treatment with 10-320 μM DHA or OA control fatty acid for 72h (Fig **1**). OA treatment did not affect the metabolic activity of any of the OC cell lines tested (Fig. S1). In contrast, DHA significantly reduced the metabolic activity of ES2 (IC_50_=30 μM) and A2780cp (IC_50_=228.7 μM) but did not affect Kuramochi and SKOV3 cells (Fig **1**); however, in crystal violet assays, viability of both Kuramochi and SKOV3 was significantly reduced by DHA with IC_50_ values of 74 μM and 254 μM, respectively (Appendix Fig **S2**).

**Figure 1.**
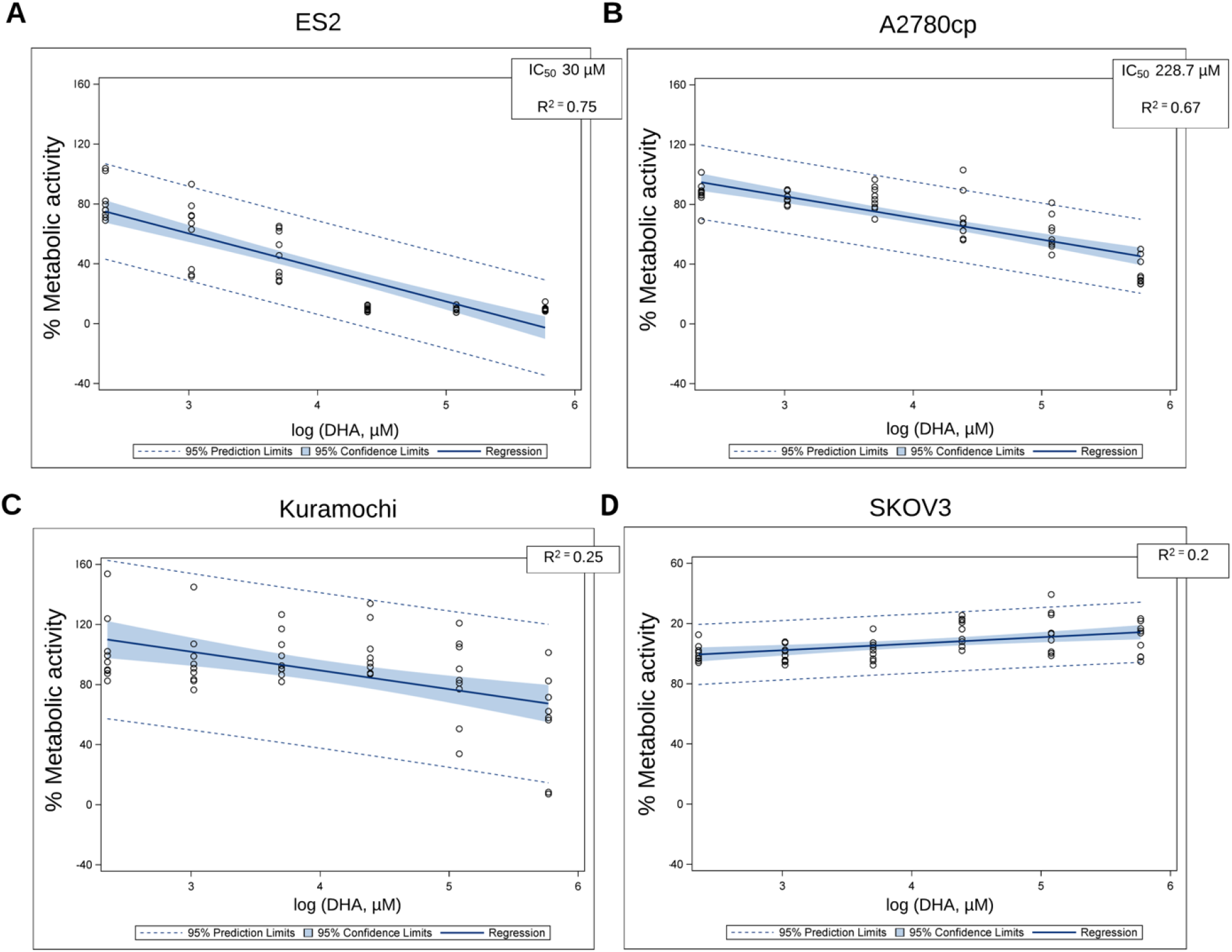
DHA dose-response linear regression curves generated from MTT assays in (A) ES2, (B) A2780cp, (C) Kuramochi and (D) SKOV3 cells. Cells were treated with DHA at 10-320 μM for 72h. The IC_50_ for DHA in ES2 and A2780cp cells was determined by fit curve linear regression in SAS Ver. 9.4.

ES2 cells treated with DHA at the IC_50_ value of 30 μM or OA at 30 μM for 48 and 72h were stained with Annexin V (AnnV) and Zombie NIR (ZNIR) followed by acquisition by flow cytometry (Fig **2A**; Appendix Fig **S3**). Percentages of early apoptotic (AnnV^+^ZNIR^−^), late apoptotic (AnnV^+^ZNIR^+^), and necrotic (AnnV^−^ZNIR^+^) cells were determined after each treatment time point. Relative to untreated or OA-treated ES2 cells, incubation with DHA resulted in markedly increased apoptosis after both time points (late apoptotic cells comprised 10.5% after 48h and 33.2% after 72h treatment with DHA compared to 3.3% and 3.5%, respectively, after treatment with OA; (Fig **2A;** Appendix Fig **S3A**), whereas treatment with DHA or OA at 160 μM for 48 and 72h did not induce apoptosis in Kuramochi cells (Fig **2B;** Appendix Fig **S3B**).

**Figure 2.**
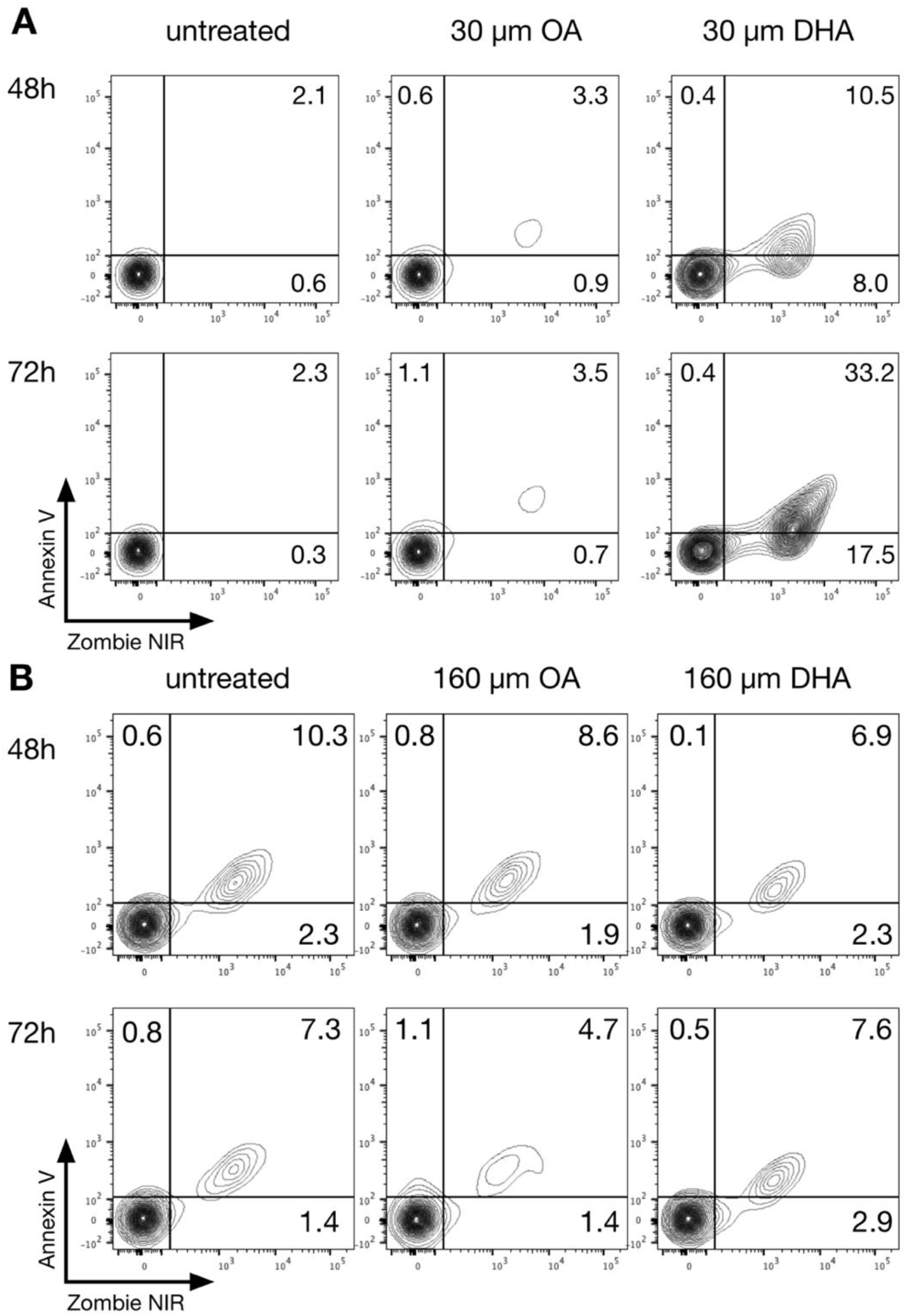
DHA induces apoptosis in ES2 but not Kuramochi OC cells. (A) ES2 cells were treated with DHA at the IC_50_ concentration (30 μM) or 30 μM OA as a control for 48 or 72h followed by flow cytometry to identify apoptotic cells. (B) Kuramochi cells were treated with 160 μM DHA or OA for 48 or 72h followed by flow cytometry. The percentage of early apoptotic [Annexin V (AnnV)^+^ Zombie NIR (ZNIR)^−^], late apoptotic (AnnV^+^ZNIR^+^) and necrotic (AnnV^−^ZNIR^+^) cells were determined at each time-point and are indicated in the quadrants.

In normal immortalized FT194 and IOSE-364 cells, 10-320 μM DHA and OA did not reduce metabolic activity after 72h treatment in MTT assay, (Fig **S4A and B**); however, some evidence of 160 μM DHA inducing apoptosis was observed in FT194 cells treated for 72h, with 21.1% late apoptotic cells compared to 2.8% in OA-treated cells (Appendix Fig **S4C and D**).

### DHA and carboplatin in combination kill ovarian cancer cells to a greater extent than either treatment alone in vitro

To study potential combinatorial effects, Kuramochi and SKOV3 cells were incubated with different concentrations of DHA and carboplatin for 72h before measuring surviving attached cells by crystal violet staining and colorimetry. We observed synergistic cell killing by DHA and carboplatin in OC cancer cell lines Kuramochi and SKOV3, as demonstrated in heat maps indicating percentage cell viability normalized to untreated cells (Fig **3A and B**). We also calculated Bliss combination indices (CIs), which indicate synergy for values below 1 (Fig **3C**). For example, treatment with 20 μM DHA and 6 μM carboplatin for 72h alone did not affect cell viability of SKOV3 cells (cell viability 97.9 and 85%, respectively); however, combining DHA and carboplatin at these concentrations resulted in 62% viability (Fig **3B**). The Bliss CI was 0.43 indicating a synergistic effect. We observed a similar trend in Kuramochi cells (Fig **3A**). Treatment with DHA and carboplatin alone at concentrations of 10 and 3 μM, respectively, resulted in cell viability of 83 and 89%. After combination of these concentrations the cell viability was decreased to 69% and Bliss CI was 0.84.

**Figure 3.**
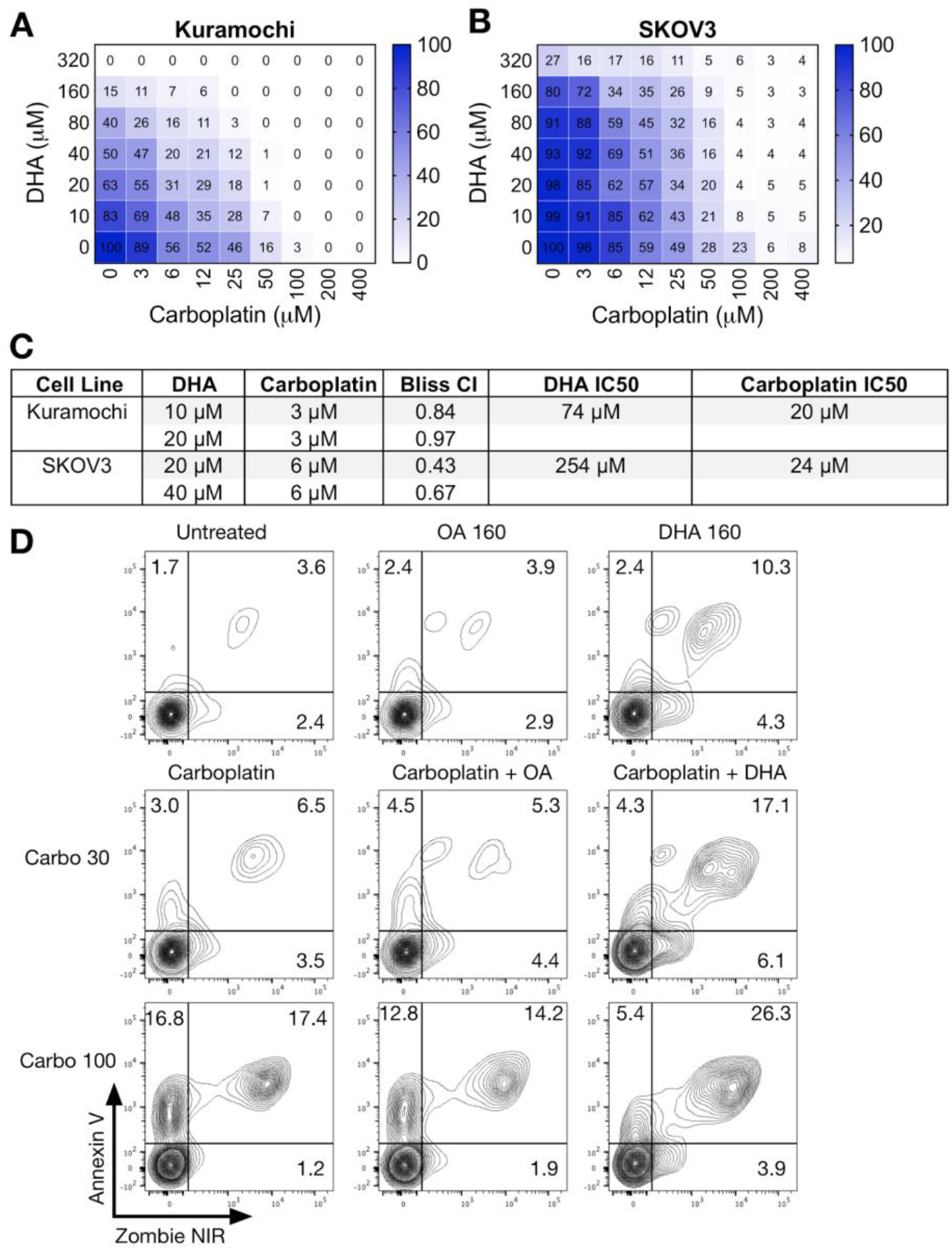
Synergistic killing of Kuramochi and SKOV3 cells by combination treatment with carboplatin and DHA. (A) Kuramochi and (B) SKOV3 cells were treated for 72h with a combination of up to 400 μM carboplatin and up to 320 μM DHA. Cell viability was assessed by crystal violet assay. Data were generated from three independent biological replicates. Color bars in heat maps indicate percentage cell viability normalized to untreated cells. (C) Table showing Bliss combination indexes (CIs) and IC_50_ for DHA and carboplatin. Bliss CI <1 indicates synergy. Heat maps show that the combination of DHA with carboplatin significantly reduces cell viability of both cell lines compared to DHA or carboplatin treatment alone. (D) The combination of DHA with carboplatin induces more apoptosis compared to carboplatin or DHA alone in Kuramochi cells. Cells were treated for 72h with 30 or 100 μM carboplatin alone, 160 μM DHA or OA alone and or a combination of 30 or 100 μM carboplatin with 160 μM OA or DHA, stained with Zombie NIR and AnnexinV, then acquired by flow cytometry.

Flow cytometric analysis indicated that 72h treatment with a combination of 160 μM DHA and 30 or 100 μM carboplatin induced more apoptosis in Kuramochi cells compared to carboplatin alone, while treatment with DHA or OA alone did not induce appreciable apoptosis (Fig **3D**). Relative to carboplatin or carboplatin/OA-treated Kuramochi cells, combination of DHA with carboplatin at both concentrations resulted in markedly increased apoptosis (late apoptotic cells comprised 17.1% and 26.3% after treatment with DHA and 30 or 100 μM carboplatin compared to 5.3% and 14.2%, respectively, after treatment with OA in combination with carboplatin.

### Incorporation of long chain polyunsaturated fatty acids in ovarian cancer whole cell phospholipids

Assessment of the relative composition (percentage) of whole cell membrane PLs revealed higher DHA and ∑N-3 PUFA content in OC cells ES2 and A2780cp treated with DHA compared to those treated with OA, although the difference did not reach significance in SKOV3 cells (Appendix Table **S2**). In ES2 and A2780cp cell lines, as 72h treatment with DHA significantly increased the proportion of DHA and ∑N-3 PUFA and decreased levels of ∑N-6 PUFA in the whole-cell PLs (p<0.05). In the OA-treated cells, the differences in fatty acid composition were not significant. In contrast, treatment with DHA and OA did not induce significant changes in fatty acid levels in whole-cell PLs of DHA-resistant SKOV3 cells.

### DHA dietary supplementation reduces tumor growth *in vivo*

To determine whether DHA improves the efficacy of chemotherapy, DHA-fed mice bearing PDX-550 tumors were treated with carboplatin or vehicle control, and tumors were extracted at an experimental endpoint of 10 mm in diameter (Fig **4A**; Appendix Fig **S5A**). Notably, carboplatin was administered IP (to mimic suboptimal treatment levels) in the first experiment and IV in the second experiment (to represent optimal delivery). DHA-fed mice exhibited a reduction in tumor growth in all experimental groups, with or without carboplatin (Fig **4B, C, D and F**; Appendix Fig **S5B and C**). DHA with IP administration of carboplatin did not significantly affect tumor growth compared to DHA supplementation alone (Fig **4E**); however, tumors from mice fed the DHA diet were sensitized to carboplatin compared to the control diet combined with carboplatin (Fig **4F**, DHA + carboplatin Slope 0.6 *versus* control + carboplatin Slope 1.6, p=0.0006). When mice received IV treatment, DHA alone had the same effect on tumor growth as carboplatin with control diet (p=0.8) or carboplatin with DHA supplementation (p=0.2, Fig. S5E, S5F). Since the endpoint was chosen based on tumor size, final tumor volumes and weights at endpoint did not significantly differ among treatments (Fig **4G, H and I**; Appendix Figs **S5G, H and I**). In the experiment testing the IP route of carboplatin administration, the overall survival (time to endpoint) of mice bearing PDX-550 tumors was longer in DHA groups with and without carboplatin treatment (median survival 32 days) compared to controls (median survival 25 days in both control diet groups). The overall survival of mice in the experiment with IV administration of carboplatin was also longer in the DHA diet group (median survival 59 days) compared to the control diet group (43.5 days); however, survival times did not differ between groups on control or DHA diets in combination with carboplatin chemotherapy (median survival 63 days for both groups) (Appendix Fig **S4J**). Histological examination demonstrated that DHA alone or in combination with carboplatin significantly increased necrotic areas of tumors compared to control tumors (Fig **5A and B**; Appendix Fig **S6A and B**). There was no difference in the Ki-67 proliferation index among different groups considering that all tumors were collected at one endpoint (Fig **5C and D**; Appendix Fig **S6C and D**).

**Figure 4.**
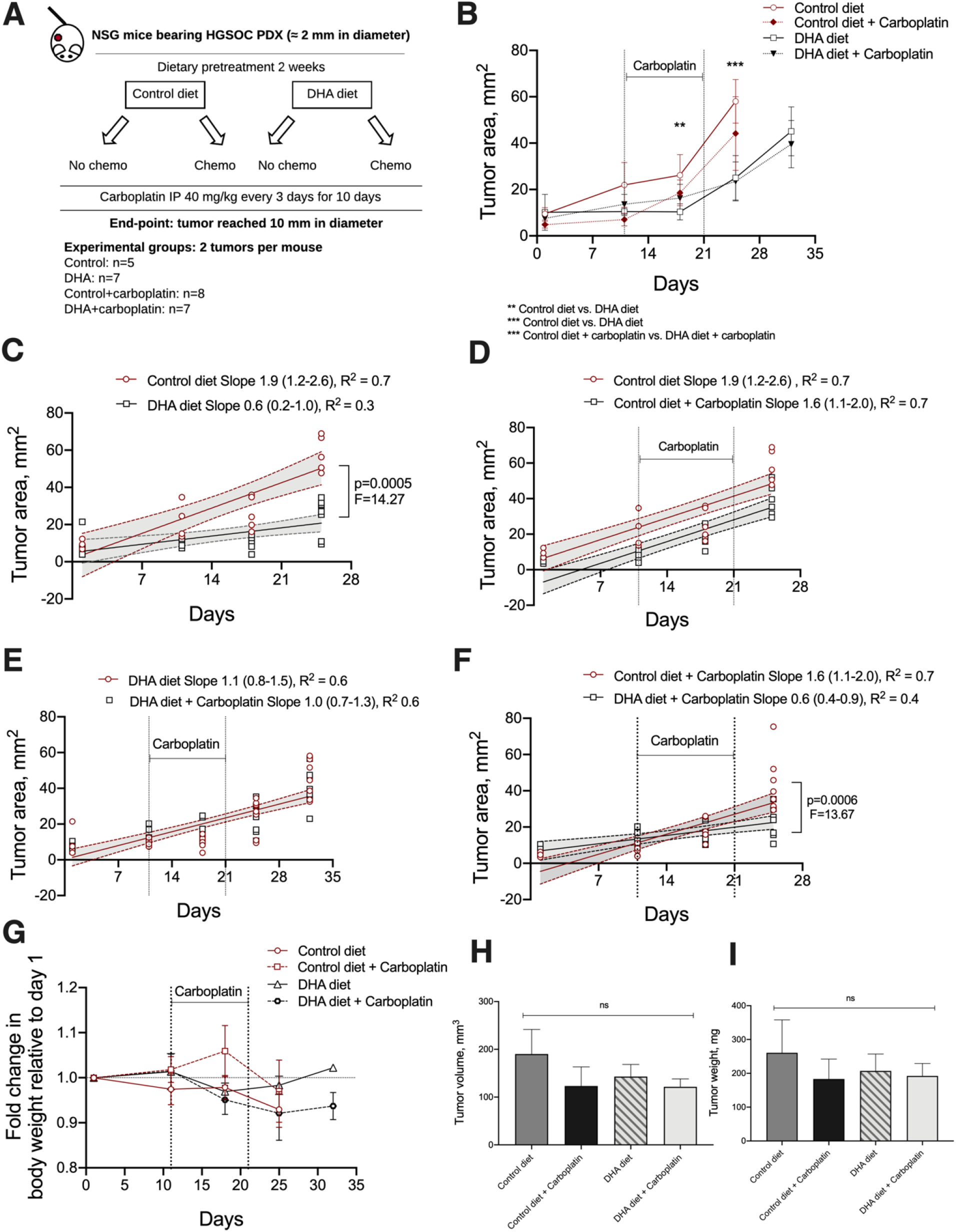
Effect of dietary DHA with or without intraperitoneal (IP) carboplatin administration on HGSOC PDX-550 growth in NSG mice. (A) Experimental design: mice were implanted subcutaneously with HGSOC PDX-550 tumor sections approximately 2 mm in diameter. Two weeks prior to commencing chemotherapy, the mice were randomized into control or DHA diet groups. They were then subsequently further randomized into one of the groups receiving carboplatin chemotherapy or vehicle control *via* IP administration. Carboplatin administration started after two weeks of dietary pre-treatment when tumors reached approximately 10-20 mm^2^. Experimental groups are defined as control diet, control diet + carboplatin, DHA diet, and DHA diet + carboplatin. The experimental endpoint was defined as the point at which the tumor reached 10 mm in diameter (~60 mm^2^). (B) Average tumor area of HGSOC PDX-550 tumor bearing mice over time. The difference between groups was determined by multiple unpaired t-tests. ** statistical difference between day 18 of DHA diet compared to control, *** statistical difference from day 25 of DHA compared to control and DHA + carboplatin compared to control + carboplatin. (C) Non-linear regression curve fit of the straight line showing significant difference in slopes of tumor growth between control diet and DHA diet groups, (D) control diet and control diet + carboplatin groups and (F) control diet + carboplatin and DHA diet + carboplatin groups. (E) Non-linear regression curve fit of the straight line showing no differences in tumor growth between DHA and DHA + carboplatin groups. Values in non-linear regression curve fit represent individual replicates with 95% Confidence Interval. (G) Fold change in body weight of mice randomized into different groups relative to day one. (H) Excised tumor volume (mm^3^) and tumor weight (I) of PDX tumor bearing mice at the endpoint. Values represent mean±SD.

**Figure 5.**
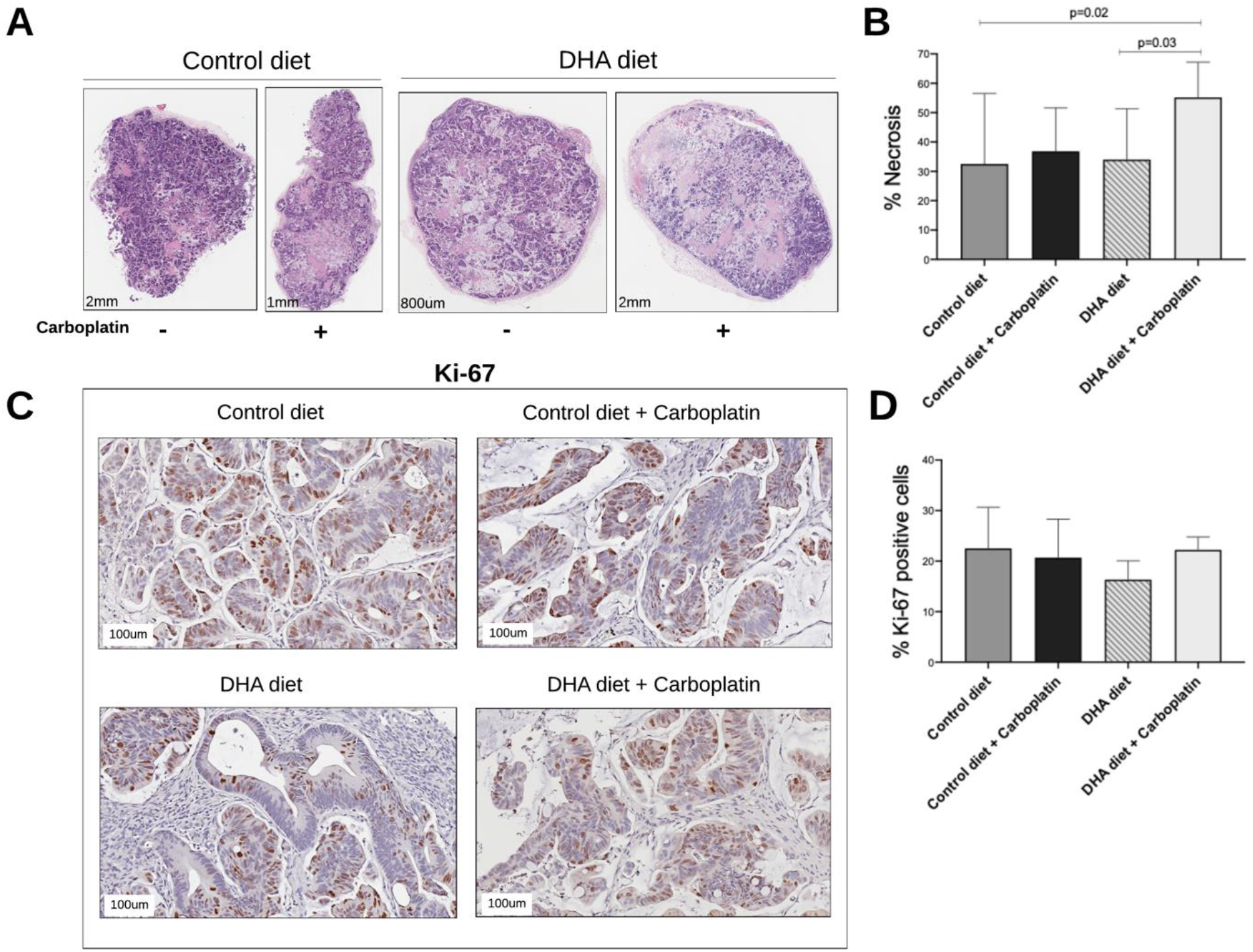
Effect of dietary DHA with or without intraperitoneal (IP) carboplatin administration on HGSOC PDX-550 tumor necrosis and cell proliferation in NSG mice. (A) Representative H&E images of extracted tumors from NSG mice randomized into control or DHA groups with or without carboplatin administration. (B) Average area of necrosis in tumor extracts from NSG mice randomized into different groups. Tumors extracted from the DHA + carboplatin group show significantly larger areas of necrosis compared to control diet and DHA diet groups (p<0.05). (C) Representative images of Ki-67 IHC staining in tumor sections. (D) Number of cells expressing the proliferation marker Ki-67 in tumor sections from NSG mice randomized into different groups. Tumors extracted from different groups do not differ in proliferation rate. Values represent mean±SD.

### Incorporation of Long Chain Polyunsaturated Fatty Acids in Tumor and Liver Phospholipids

We further assessed the content of LCPUFA – arachidonic acid (ARA), eicosapentaenoic acid (EPA), docosapentaenoic acid (DPA) and DHA, and a total content of saturated (∑SFA), monosaturated (∑MUFA), and polysaturated fatty acids (∑PUFA), ∑N-3 PUFA and ∑N-6 PUFA – in tumor and liver PLs for all treatment groups (Appendix Table **S3**). The DHA-enriched diet significantly increased the percentage of DHA in the plasma membrane of the tumor tissues from 2.96% to 4.62% and total ∑N-3 PUFA from 4.15% to 6.64% (p<0.05) versus the control diet but decreased the concentration of ARA and ∑N-6 PUFA. We observed the same trend in the DHA versus control diet groups treated with carboplatin. This was also true for analyses of whole-cell PLs in livers.

## DISCUSSION

In this study, we used four different cell lines and a HGSOC PDX model to illuminate an anti-tumorigenic role for DHA in ovarian cancers, irrespective of intra-or inter-tumoral heterogeneity. This finding supports previous studies, which have shown that DHA can inhibit cancer cell proliferation and can elicit selective cytotoxicity in cell lines derived from colon, gastric, pancreatic, lung, breast and ovarian cancers (Song & Kim, 2016; Wan *et al*, 2016).

Collectively, our study adds to a growing body of literature suggesting that DHA may be a desirable addition to adjuvant cancer therapies.

Our *in vitro* findings demonstrate that DHA shows a variable inhibitory effect on cell viability and apoptosis in OC cell lines and normal immortalized cells with little or no toxicity to normal cells. Among all cell lines tested, ES2 cells were the most sensitive to DHA in MTT assays with an IC_50_ of 30 μM. The significant inhibitory effect of DHA on ES2 cells in our experiments may be due to the fact that these cells express wild type p53. Indeed, a recent study showed that DHA significantly suppresses growth in TOV-21G cells concomitant with increased PPARγ and p53 expression (Wan *et al*, 2016). Interestingly, DHA did not affect Kuramochi and SKOV3 cells in MTT assays but inhibited cell viability in crystal violet assays without causing significant apoptosis, suggesting alternative mechanisms of action in these p53 mutant cell lines. While DHA did not profoundly affect SKOV3 and Kuramochi viability as a monotherapy, we found that the combination of DHA and carboplatin worked synergistically to increase cell death in both cell lines. This is in agreement with previous study, which showed that DHA significantly enhanced the cytotoxicity of cisplatin in SKOV3 and OVCAR3 cells (Zajdel *et al*, 2018).

To extend our *in vitro* findings, we used a HGSOC PDX model to investigate how DHA supplementation affects OC growth in combination with cytotoxic chemotherapy administered IP or IV. DHA-fed mice exhibited a significant reduction in tumor growth in all experimental groups, even without carboplatin. This reduction in growth was accompanied by increased levels of necrosis. While DHA did not improve the effects of carboplatin delivered IV, the DHA-enriched diet increased sensitivity of PDX tumors to carboplatin when administered IP. These results corroborate other preclinical studies showing that DHA can inhibit tumor growth, metastasis and drug resistance (Newell *et al*, 2019; Song & Kim, 2016, Merendino *et al*, 2013).

Our findings also illustrate the safety of DHA. DHA was not cytotoxic against normal ovarian surface epithelial cells or fallopian epithelial cells, and mice tolerated the DHA diet well as their body weights did not significantly change over the duration of the experiments. This is in accordance with previous studies wherein DHA was cytotoxic to breast cancer cells but did not affect the viability of human peripheral blood mononuclear or normal mammary epithelial cells (Pizato *et al*, 2018). Collectively, this, together with clinical trials which have administered DHA to patients (Morland *et al*, 2016), suggests that DHA would have minimal expected toxicity in a clinical setting.

DHA incorporates into plasma membranes, altering microdomain organization and affecting essential cell signaling events (Corsetto *et al*, 2011; Sawyer & Field, 2010; Ibarguren *et al*, 2014) We confirmed that DHA, but not OA, successfully incorporated into the plasma membranes of ES2 and A2780cp cells. However, neither DHA nor OA induced significant changes in fatty acid levels in whole-cell PLs of DHA-resistant SKOV3 cells. Indeed, it appears that sensitivity to DHA is dependent on its efficient incorporation into plasma membranes. To support this notion, we found that plasma membranes in tumours derived from mice fed the DHA-enriched diet had an increase in DHA (2.96% to 4.62%) as compared to those in tumors extracted from mice fed a control diet. Total N-3 PUFA also increased from 4.15% to 6.64% and the concentrations of ARA and total N-6 PUFA were reduced. The inverse correlation between N-6 and N-3 PUFA in the DHA diet group alone or in combination with carboplatin indicates that membrane incorporation of N-3 PUFA partially replaces ARA, as has been shown in other studies (D’Eliseo & Velotti, 2016). Finally, similar results were obtained using whole-cell PLs derived from livers, suggesting that tumors take up DHA as effectively as normal tissues.

The incorporation of DHA into lipid rafts affects the content and function of myriad transmembrane proteins (including growth factor receptors, and drug transporters), which may collectively regulate pro-tumorigenic processes (Corsetto *et al*, 2017). Moreover, DHA has been shown to promote apoptosis and to induce cell cycle arrest in various cancer cell types, by regulating several putative pathways (Song & Kim, 2016; Newell *et al*, 2017). Given the pleiotropic mechanisms associated with DHA, as well as our focus on testing the clinical potential of DHA for the treatment of OC, this study did not interrogate the mechanisms by which DHA promotes cytotoxicity in OC cells, beyond demonstrating the induction of apoptosis. Future studies should use a systems biology approach in order to better understand how numerous mechanisms coalesce to effectuate the anti-cancer effects of DHA.

Several recent clinical trials have shown the association between adjuvant N-3 PUFA supplementation and improved chemotherapy tolerability and patient outcomes. These trials were performed in different cancer populations including pancreatic, gastrointestinal, breast, colorectal and lung cancers using a variety of chemotherapy regimens (Bougnoux *et al*, 2009). These studies reported that ≤6 g of encapsulated oil per day was well tolerated by patients and no serious adverse events were reported related to the study interventions. A double-blinded, phase II, randomised controlled trial of 52 women prescribed neoadjuvant chemotherapy for breast cancer is currently testing if DHA supplementation at dose of 5g per day for 12-18 weeks enhances efficacy of chemotherapy (NCT03831178). If the results from this study are positive, they will provide further rationale for supplementation with DHA during neoadjuvant chemotherapy treatment for breast cancer (Newell *et al*, 2019). To date, there have been no clinical trials testing the efficacy of PUFA supplementation on clinical outcomes in OC. However, an active randomized clinical trial pilot is being conducted to determine whether omega-3 fatty acids can treat pain in patients with breast or ovarian cancer receiving paclitaxel (NCT01821833).

Collectively, our study, together with the extant literature, suggests that treatment with N-3 LCPUFA could improve sensitivity to chemotherapy in patients with chemoresistant recurrence, without inducing significant adverse events. By increasing our understanding of how a simple nutritional intervention such as DHA supplementation can improve conventional drug treatment, our novel study provides a strong rationale to move directly to clinical trials, to determine whether DHA increases the efficacy of cytotoxic chemotherapy for treatment of OC.

## MATERIALS AND METHODS

### Cell Lines

ES2 (ATCC, Cat # CRL-1978) and SKOV3 (ATCC, Cat # HTB-77) cells were obtained from American Type Culture Collection (ATCC) and cultured in McCoy’s 5A medium (MilliporeSigma, Cat # M8403). Kuramochi (JCRB Cat # JCRB0098) cells were obtained from Japanese Collection of Research Bioresources Cell Bank (JCRB) and cultured in RPMI1640 (MilliporeSigma, Cat # R7388). Immortalized ovarian surface epithelial cells IOSE-364 (RRID:CVCL_5540) were obtained from Canadian Ovarian Tissue Bank, University of British Columbia and cultured in DMEM/F12 medium (MilliporeSigma, Cat # DF-041)). A2780cp (NCBI_Iran Cat# C454) cells were kindly provided by Dr. Benjamin Tsang (Ottawa Hospital Research Institute) and cultured in DMEM/F12 medium (MilliporeSigma, Cat # DF-041). All medium was supplemented with 10% fetal bovine serum (FBS, Thermo Fisher, Cat # A4766801) and cells were cultured at 37°C in 5% CO_2_. Immortalized fallopian tube secretory epithelial cells FT194 (RRID:CVCL_UH58) (Karst & Drapkin, 2012) were kindly provided by Dr. Drapkin’s laboratory (University of Pennsylvania) and cultured in DMEM/F12 (Thermo Fisher, Cat # 11320033) supplemented with 2% Ultroser G (Pall France, Cat # 15950-017). All cells were used for up to 8 passages after thawing. All OC cancer cells were authenticated at the Centre for Applied Genomics, Sick Kids Research Institute. Toronto, ON, Canada (project ID LIU7072; 13 June 2018).

### Chemotherapeutic Drugs and Preparation of Bovine Serum Albumin (BSA) Conjugated Fatty Acids

Carboplatin used for *in vitro* and *in vivo* studies was purchased from MilliporeSigma (Cat # 1096407) and Cayman Chemical Company (Cat # 41575-94-4), respectively. Fatty acids were purchased from NuChek Prep, Inc. (Elysian, MN). DHA and oleic acid (OA) were prepared according to the published protocol (Ewaschuk *et al*, 2012). Fatty acids were aliquoted and stored at −20°C until further use.

### Cell Proliferation and Viability Assays

All experiments in this paper, unless otherwise indicated, were performed in triplicate (3 technical and 3 biological replicates). Cells were seeded in 96-well plates at 5000 cells per well, allowed to adhere overnight, then treated with increasing concentrations of BSA-conjugated DHA or oleic acid (OA) (10-320 μM) or carboplatin (3.1-400 μM) for 72h using the Cell Proliferation Kit I (MTT, MilliporeSigma, Cat # 11465007001) according to the manufacturer’s recommendations. The relative metabolic activity (%) was expressed as a percentage relative to untreated cells. The IC_50_ of DHA in MTT assay was determined by fit curve linear regression in SAS Ver. 9.4.

Crystal violet assays were performed to determine cell viability by detecting adherence of cells following carboplatin and DHA/OA treatment. The protocol was adapted from (Feoktistova *et al*, 2016). Relative cell viability (%) was expressed as a percentage relative to untreated cells.

The IC_50_ of carboplatin and DHA in OC cells in MTT and crystal violet assays was determined by fit curve non-linear regression using GraphPad Prism Ver. 7.0. For Bliss drugs’ independence, cells were treated as described above alone or in combination and Bliss combination indices were calculated as previously described (Foucquier & Guedj, 2015). Heat maps were generated to illustrate the percentage cell viability normalized to untreated cells after treatment with different dose combinations of DHA and carboplatin.

### Phospholipid Fatty Acid Composition Analysis

Phospholipids (PLs) from experimentally treated whole cell lysates as well as ground tumor and liver tissues were extracted by a modified Folch procedure as previously described (Field *et al*, 1988; Folch *et al*, 1957). Total PLs were separated by thin layer chromatography and the fatty acid content was measured by gas-liquid chromatography (GLC 7890A, Agilent Technologies) on a CP-Sil 88 100 m column (Agilent Technologies) (Cruz-Hernandez *et al*, 2013).

### Detection of Apoptosis

ES2 cells were treated with the IC_50_ of DHA (30 μM) and corresponding concentration of OA for 24, 48 and 72h. Kuramochi and FT194 cells were treated with 160 μM of DHA and OA for 24, 48 and 78h. For evaluating combination effects, Kuramochi cells were cultured with 30 and 100 μM carboplatin combined with 160 μM DHA or OA for 72h. Cell supernatants were collected and combined with trypsinized cells to ensure that both live and dead cells were analyzed in the assay. After centrifugation and washing with 1 x phosphate buffered saline (PBS), cells were stained with Zombie NIR fixable viability dye (BioLegend, Cat # 423106) following the manufacturer’s instructions. Cells were washed with 1 x Annexin V (AnnV) binding buffer (BioLegend, Cat # 422201) and then stained with AnnV BV510 (BioLegend, Cat # 640937, 1:20) on ice for 15 min in the dark. Cells were then washed and re-suspended in 200 μl 2% paraformaldehyde (MilliporeSigma, Cat # 1004960700) in AnnV binding buffer and stored at 4°C until analysis (within one week). Samples were acquired on a FACS Canto II (Becton Dickinson) flow cytometer. Forward and side scatter gating was done to exclude debris; fluorescence minus one single-stained controls were used for quadrant gates. Analysis was performed using FlowJo© software (Tree Star Inc, Version 10.5.3).

### Experimental Diets and *In Vivo* Studies

A nutritionally complete basal mix diet from Teklad (TD.84172, Harlan Laboratories, Madison, WI) was supplemented with 20% w/w fat. The composition of fatty acids in the diets (Table S1) was achieved by blending oils to obtain a DHA content of 3.9% w/w of total fat (DHASCO^TM^, DSM, Columbia, MD). This amount of dietary DHA was selected to achieve a plasma PLs DHA concentration of > 5% w/w of total fatty acids as this concentration has been associated with improved outcomes in patients with metastatic breast cancer receiving chemotherapy in a phase II clinical trial (Bougnoux *et al*, 2009).

Animal experiments were approved by the University of Alberta Animal Policy and Welfare Committee (AUP00002496) and were in accordance with the Canadian Council on Animal Care guidelines. The PDX OVXF-550 (high-grade serous adenocarcinoma – HGSOC) model was obtained from Charles River Oncotest^TM^ (Germany). PDX OVXF-550 has a *TP53* mutation and is poorly differentiated with high vascularization (Online resource: https://compendium.criver.com/compendium2/model?model.id=10067&query.key=525c0877993f601965cb0a1d8ca5d45c). Up to two PDX tumor sections approximately 2 mm in diameter were implanted subcutaneously into 6-8-week-old female NOD.Cg-Prkdc^SCID^Il2rg t^m1Wjl^/SzJ (NSG) mice (The Jackson Laboratory, Stock # 005557). Once tumors reached approximately 5-10 mm^2^ (approximately 3 weeks after implantation), mice were randomized into two diet groups – 0% DHA (control) or 3.9% w/w DHA (DHA) – and fed these diets for two weeks before being further randomized for carboplatin administration by intraperitoneal (IP) or intravenous (IV) routes. In the first experiment, mice received 40 mg/kg carboplatin or vehicle IP every 3 days for 10 days (total 4 injections, Fig. 4A). In the second experiment, mice received 70 mg/kg carboplatin by tail vein once a week for three weeks (total 3 injections, Figure S5). Experimental groups were defined as: control diet, control diet + carboplatin, DHA diet, and DHA diet + carboplatin. Body weight and tumor size were measured twice a week (by caliper) and the area of tumors was calculated with the equation (area of ellipse): tumor area (mm^2^) = π x length (mm) x width (mm)/4.

The experimental endpoint was attained when a tumor reached 10 mm in diameter (~60mm^2^), at which time mice were euthanized, and tumors were carefully excised and weighed; one piece was fixed in formalin and another stored at −80°C for whole PLs analysis. Livers were also excised and stored at −80°C for whole PLs analysis. Excised tumors were measured by caliper and tumor volume was calculated with the equation: tumor volume (mm^3^) = π x length (mm) x width (mm) x height (mm)/6.

### Analysis of Necrosis and Immunohistochemistry

After staining with H&E, the entire cross-sectional area of all the slides was scanned on the Aperio Digital Pathology slide scanner (Leica Biosystems). Histological features were evaluated using Image Scope software (Ver. 12.3, Leica Biosystems). To quantify the abundance of the necrosis per tumor cross-sectional area, the whole slide image was converted to TIFF format and analysed in ImageJ (Ver.1.53). The area of necrosis was measured using a first round of thresholding against background pixel intensity compared to the intact tumor tissue, after which a second round of thresholding against necrotic regions was performed. The area of necrosis was calculated as a percentage over the total tumor area.

Formalin-fixed, paraffin-embedded tumor sections (two serial sections per sample) were used for immunohistochemistry (IHC) analysis of Ki-67 expression. Briefly, following antigen retrieval by boiling for 20 min in target retrieval solution (pH 6.0, Agilent Dako, Cat # 169984-2) and blocking steps, tumor sections were incubated with mouse anti-human Ki-67 antibody (sc-23900, Santa Cruz Biotechnology, 1:100) at 4°C overnight followed by incubation with Dako EnVision+System-HRP Labelled Polymer Anti-Mouse (Aglient Dako, Cat # K400111-2) for 60 min at room temperature. Immunostaining was detected using 3,3’-diaminobenzidine (Agilent Dako, Cat # N-1939) peroxidase chromogen substrate. Sections were counterstained with hematoxylin (Vector Laboratories, Inc, Cat # H-3401). The number of Ki-67-positively stained cells was scored in 900-1100 tumor cells from 10-15 random fields (magnification x100) by two independent investigators blinded to the treatment groups. The percentage of positive immunolabeled cells over the total number of tumor cells was calculated.

## Data Analysis

The DHA dose-response in OC cell lines was analysed by fit curve linear regression in SAS Ver.

9.4. by biostatistician Maryna Yaskina (Women & Children’s Health Research Institute). All other analyses were conducted using GraphPad Prism Ver. 7.0. The Student’s t-test was employed for comparison of a single condition to the control with paired two-sample t-tests where appropriate. Comparison of multiple conditions was done by t-tests with correction for multiple comparisons using the Holm-Sidak method. One-way ANOVAs for all pairwise comparisons with the Bonferroni and Holm post-hoc test were employed to detect statistically significant differences among multiple groups. P values < 0.05 were considered statistically significant. The IC_50_ of carboplatin and DHA in OC cells was determined by fit curve non-linear regression. Statistical differences in tumor areas were calculated with non-linear regression curve fit of the straight line by extra sum-of-squares F test and comparing the slopes.

## ACKNOWLEDGMENTS

We thank Dr. Drapkin for the fallopian tube cells. Flow cytometry was performed in the University of Alberta Faculty of Medicine and Dentistry (FoMD) Flow Cytometry Facility; this facility has received financial support from FoMD and the Canadian Foundation for Innovation awards to contributing investigators.

## Authors Contributions

OB and BH designed the experiments. OB, BH, ID, MN, ABB, RCM, JL, GMS performed the experiments. JS provided PDX tumors. GMS, AMG, CJF, LMP provided scientific input. OB and BH analyses the data. OB prepared the manuscript. LMP provided funding acquisition, supervision, review and editing

## The Paper Explained

### Problem

Despite recent advances in diagnosis and treatment, ovarian cancer (OC) is the most lethal gynecological malignancy and improving the efficacy of chemotherapy is of great interest. This study increases our understanding of how dietary intervention with docosahexaenoic acid (DHA) – a supplement proven safe for human consumption – enhances the anti-cancer effects of conventional chemotherapy.

### Results

Our study demonstrated synergistic cell killing by DHA and carboplatin in OC cell lines. Furthermore, DHA supplementation alone and in combination with carboplatin significantly reduced OC growth in a high-grade serous OC patient-derived xenograft mouse model. Carboplatin administered intraperitoneally significantly reduced tumor growth in DHA-fed mice compared to mice on the control diet. Intravenous carboplatin administration in combination with DHA reduced tumor growth similarly to carboplatin or DHA monotherapies. The DHA-induced reduction in tumor growth in this model was associated with increased tumor necrosis and improved survival.

### Impact

Patients are receptive to nutritional interventions. As such, our findings provide a strong rationale to move to clinical trials that will determine whether DHA supplementation enhances the efficacy and tolerance of cytotoxic chemotherapy in patients with OC.

---

**Supplementary Table S1.** Control *versus* DHA-enriched experimental diets formulation (W/W %)

| **Oil** | **Control diet** | **DHA diet** |
| --- | --- | --- |
| 12:0 | 0.00 | 0.34 |
| 14:0 | 1.13 | 2.19 |
| 16:0 | 22.87 | 21.08 |
| 16:1 n-9 | 1.76 | 1.80 |
| 18:0 | 13.75 | 12.39 |
| 18:1 n-9 | 36.37 | 38.66 |
| 18:1 c11 | 0.23 | 0.42 |
| 18:2 n-6 | 21.01 | 15.15 |
| 20:0 | 0.01 | 0.02 |
| 18:3 n-3 | 2.38 | 3.37 |
| 20:2 n-6 | 0.01 | 0.01 |
| 20:3 n-6 | 0.03 | 0.04 |
| 20:4 n-6 | 0.42 | 0.42 |
| 20:5 n-3 | 0.00 | 0.00 |
| 24:0 | 0.01 | 0.04 |
| 22:5 n-3 | 0.00 | 0.04 |
| 22:6 n-3 | 0.00 | 3.83 |
| ΣSFA | 37.77 | 36.05 |
| ΣPUFA | 23.85 | 22.85 |
| ΣMUFA | 38.35 | 40.88 |
| ΣN-6 | 21.47 | 15.61 |
| ΣN-3 | 2.38 | 7.24 |
| N-6/N-3 | 9.02 | 2.16 |
| P/S | 0.63 | 0.63 |

Abbreviations: MUFA, monounsaturated fatty acids; PUFA, polyunsaturated fatty acids; SFA, saturated fatty acids; P/S, polyunsaturated to saturated fatty acid ratio.

**Supplementary Table S2.**
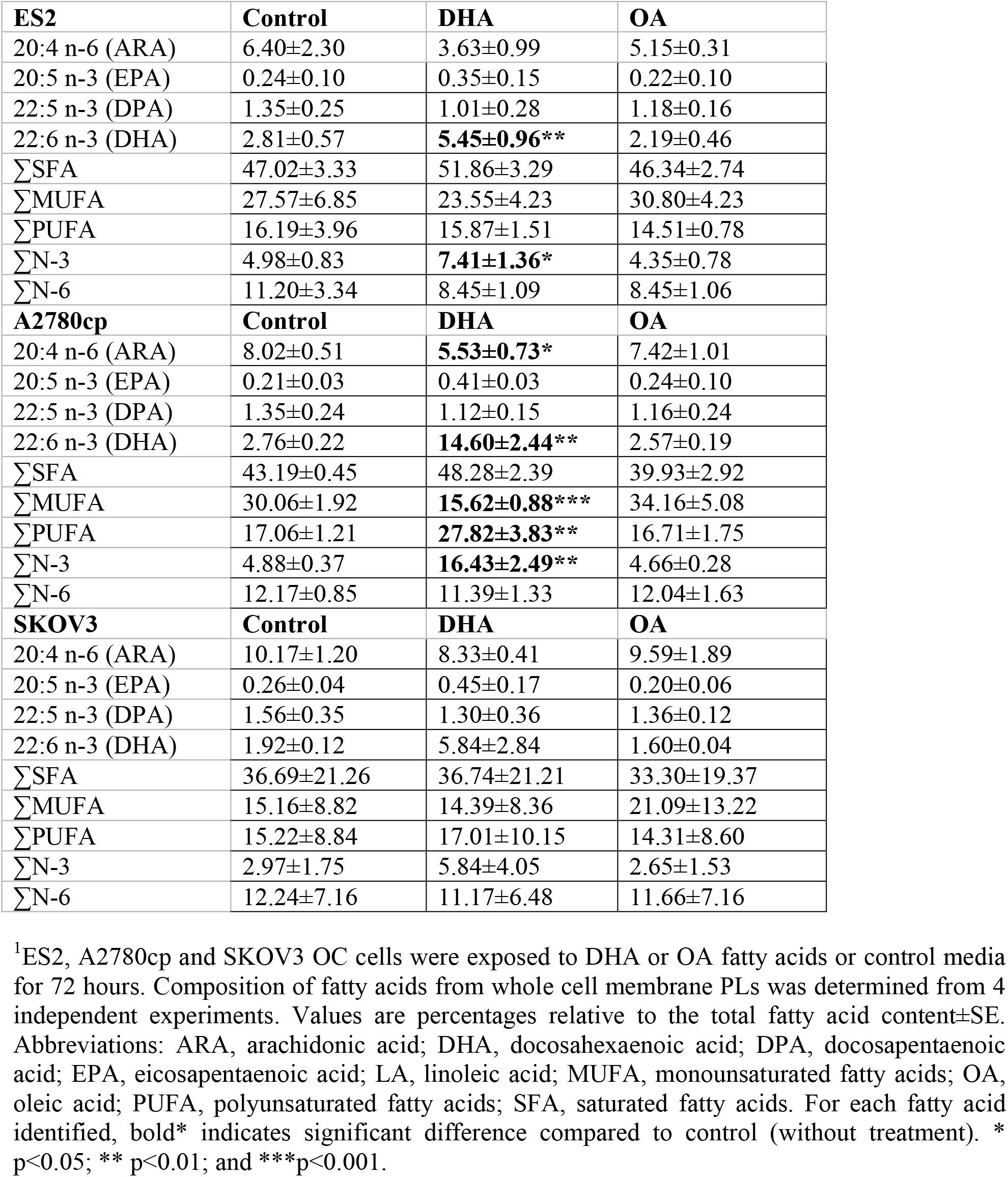
Long chain polyunsaturated fatty acid (LCPUFA) composition (%) of whole cell membrane phospholipids of ES2, A2780cp and SKOV3 OC cells^1^

**Supplementary Table S3.** Long chain polyunsaturated fatty acid (LCPUFA) composition (%) of whole cell membrane phospholipids in tumors and livers excised from NSG mice implanted with HGSOC PDX-550 tumors^1^ and treated with IP carboplatin administration (Experiment 1).

| **Tumors** | **Control** | **Control + carboplatin** | **DHA** | **DHA + carboplatin** |
| --- | --- | --- | --- | --- |
| 20:4 n-6 (ARA) | 12.48±1.14 | 12.80±1.78 | **8.13±1.25\*** | **9.33±0.52\*** |
| 20:5 n-3 (EPA) | 0.15±0.00 | 0.19±0.03 | 0.82±0.20 | 0.99±0.15 |
| 22:5 n-3 (DPA) | 0.20±0.07 | 0.19±0.06 | 0.25±0.11 | 0.23±0.06 |
| 22:6 n-3 (DHA) | 2.96±0.54 | 3.07±0.94 | **4.62±1.38\*** | **4.80±1.18\*** |
| ΣSFA | 49.63±7.62 | 51.72±8.49 | 54.36±7.96 | 53.11±8.45 |
| ΣMUFA | 14.73±8.00 | 12.51±7.59 | 12.28±6.68 | 11.27±8.24 |
| ΣPUFA | 34.46±2.66 | 34.51±3.39 | 32.06±3.79 | 34.50±2.05 |
| ΣN-3 | 4.15±0.42 | 4.28±0.83 | **6.64±1.55\*** | **6.94±1.22\*** |
| ΣN-6 | 30.30±2.43 | 30.22±2.74 | **25.41±2.42\*\*** | 27.55±1.47 |
| **Livers** | **Control** | **Control + carboplatin** | **DHA** | **DHA + carboplatin** |
| 20:4 n-6 (ARA) | 19.10±1.17 | 17.90±1.81 | **11.99±0.27\*** | **13.35±2.81\*** |
| 20:5 n-3 (EPA) | 0.09±0.03 | 0.12±0.01 | **1.48±0.08\*\*\*** | **1.32±0.17\*\*\*** |
| 22:5 n-3 (DPA) | 0.39±0.03 | 0.39±0.02 | 0.39±0.05 | 0.43±0.09 |
| 22:6 n-3 (DHA) | 11.42±1.64 | 12.27±0.92 | **17.31±0.55\*** | **17.07±2.88\*** |
| ΣSFA | 41.20±1.37 | 40.55±1.25 | 43.32±0.61 | 43.12±0.62 |
| ΣMUFA | 12.05±3.73 | 12.28±2.84 | 9.52±0.75 | 10.09±0.73 |
| ΣPUFA | 46.50±2.32 | 46.91±1.84 | 47.06±0.30 | 46.66±0.52 |
| ΣN-3 | 12.26±1.73 | 13.12±0.92 | **19.48±0.57\*** | **19.11±2.79\*** |
| ΣN-6 | 34.23±1.58 | 33.79±2.74 | **27.57±0.46\*\*** | **27.54±2.82\*** |

<sup>1</sup>NSG mice with HGSOC PDX-550 tumors were maintained on a 20% (± 3.8% DHA) w/w diet for 2 weeks before and during IP carboplatin administration. Values are percentages relative to the total fatty acid content ± SE. Within the rows, the values with different superscripts are significantly different (p<0.05). Abbreviations: ARA, arachidonic acid; DHA, docosahexaenoic acid; DPA, docosapentaenoic acid; EPA, eicosapentaenoic acid; MUFA, monounsaturated fatty acids; PUFA, polyunsaturated fatty acids; SFA, saturated fatty acids.

**Supplementary Figure S1.**
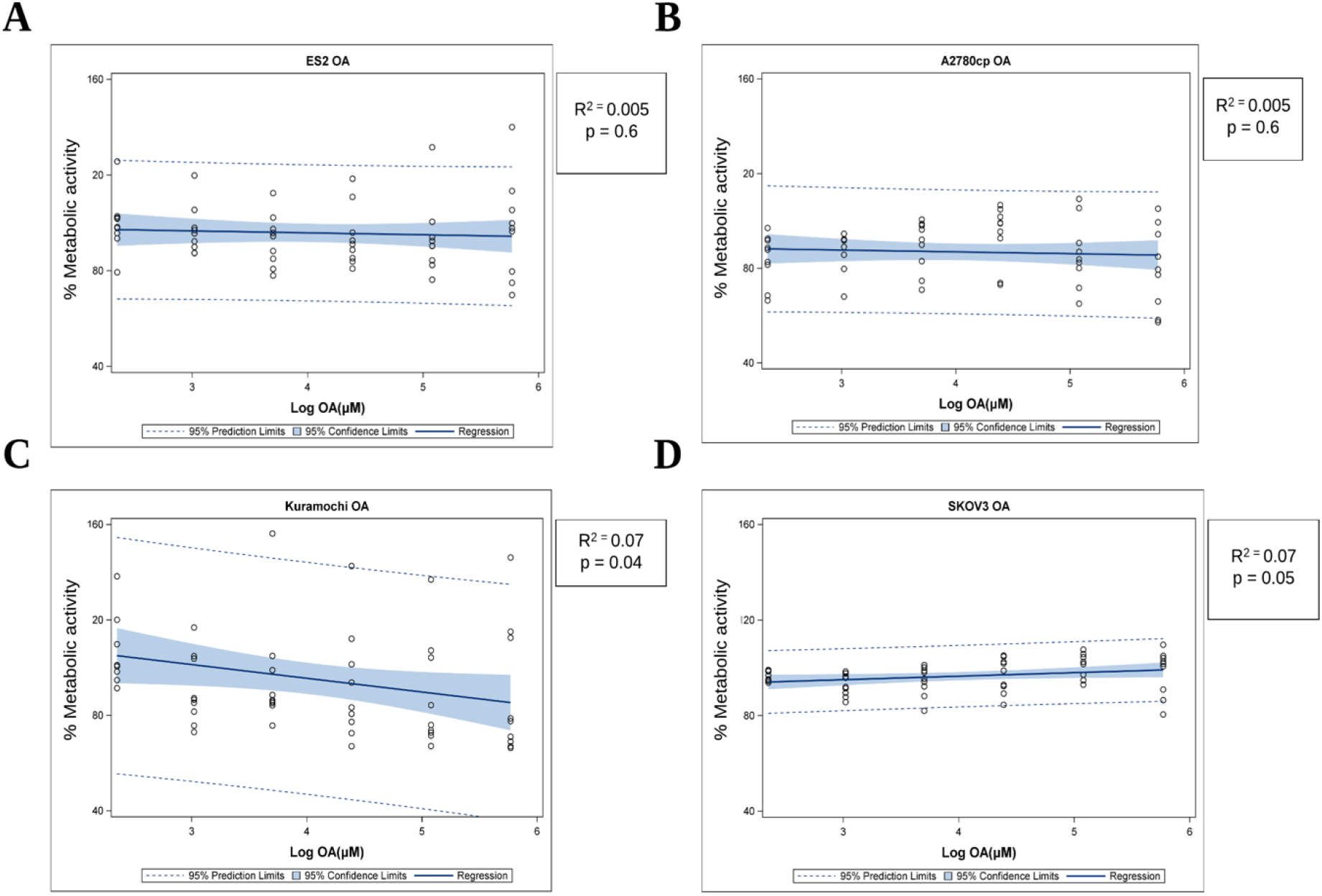
OA dose-response linear regression curves generated from MTT assays showing that OA does not affect metabolic activity of OC cells. Cells were treated with 10-320 μM OA for 72 hours. Fit curve linear regression was performed in SAS Ver. 9.4

**Supplementary Figure S2.**
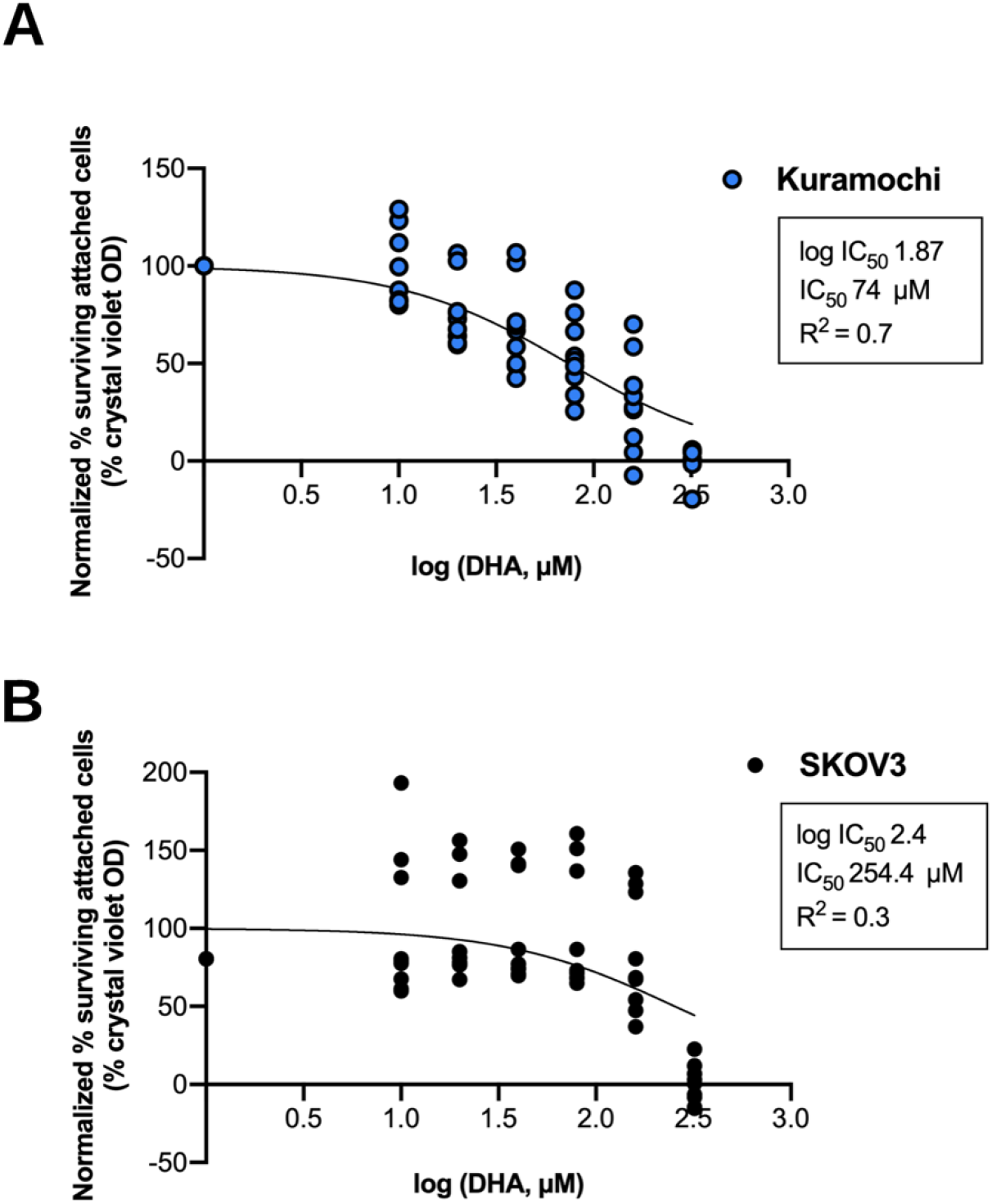
DHA reduces cell viability of (A) Kuramochi and (B) SKOV3 cells in crystal violet cell viability assays with IC_50_ of 74 μM for Kuramochi cells and 254 μM for SKOV3 cells. The IC_50_ for DHA was determined by fit curve non-linear regression. Values represent independent replicates with 95% CI.

**Supplementary Figure S3.**
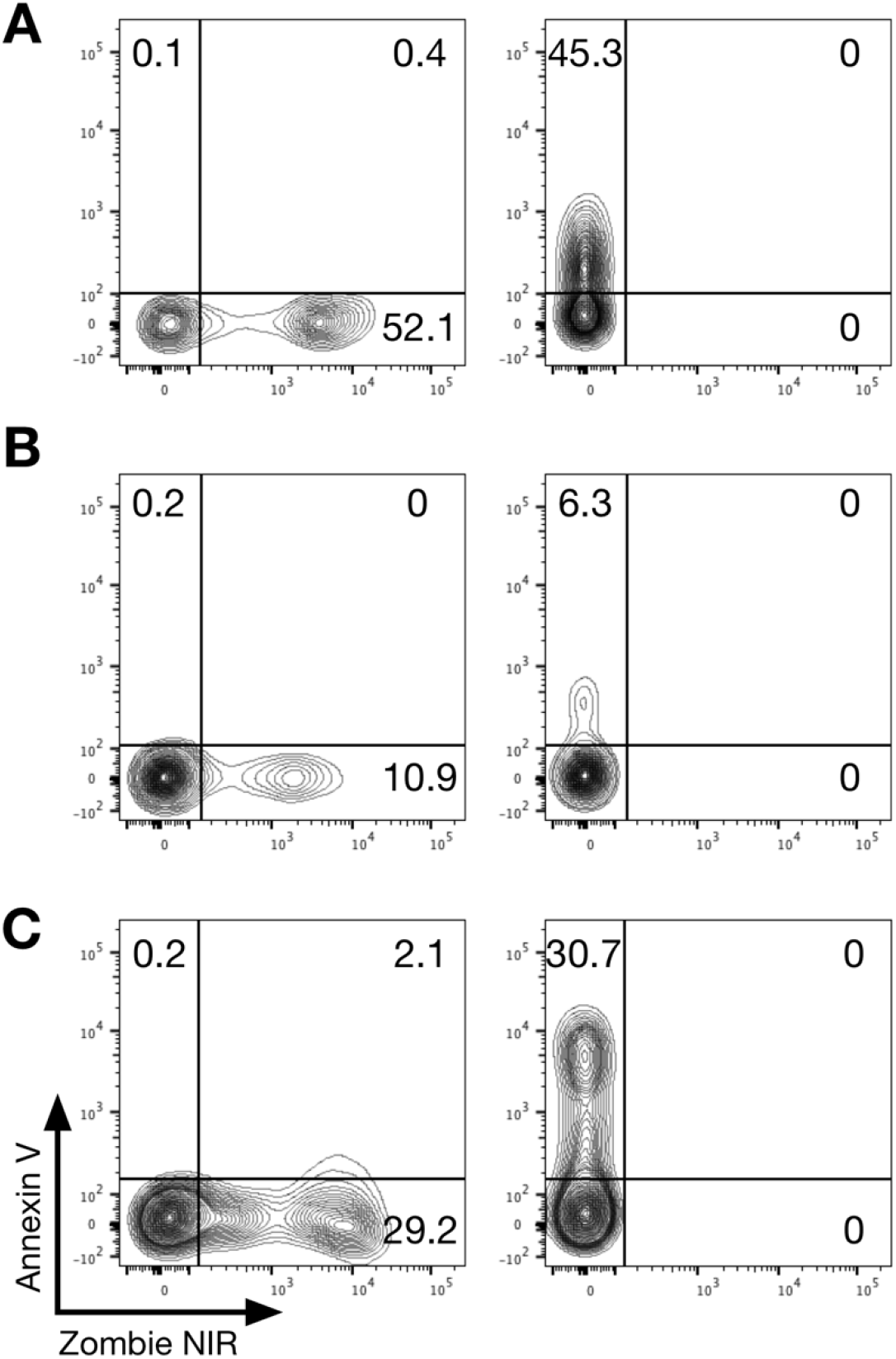
Fluorescence minus one (FMO) controls used to set quadrant gates for flow cytometry experiments shown in (A) Figure 2A, (B) Figure 2B and (C) Figure 3C.

**Supplementary Figure S4.**
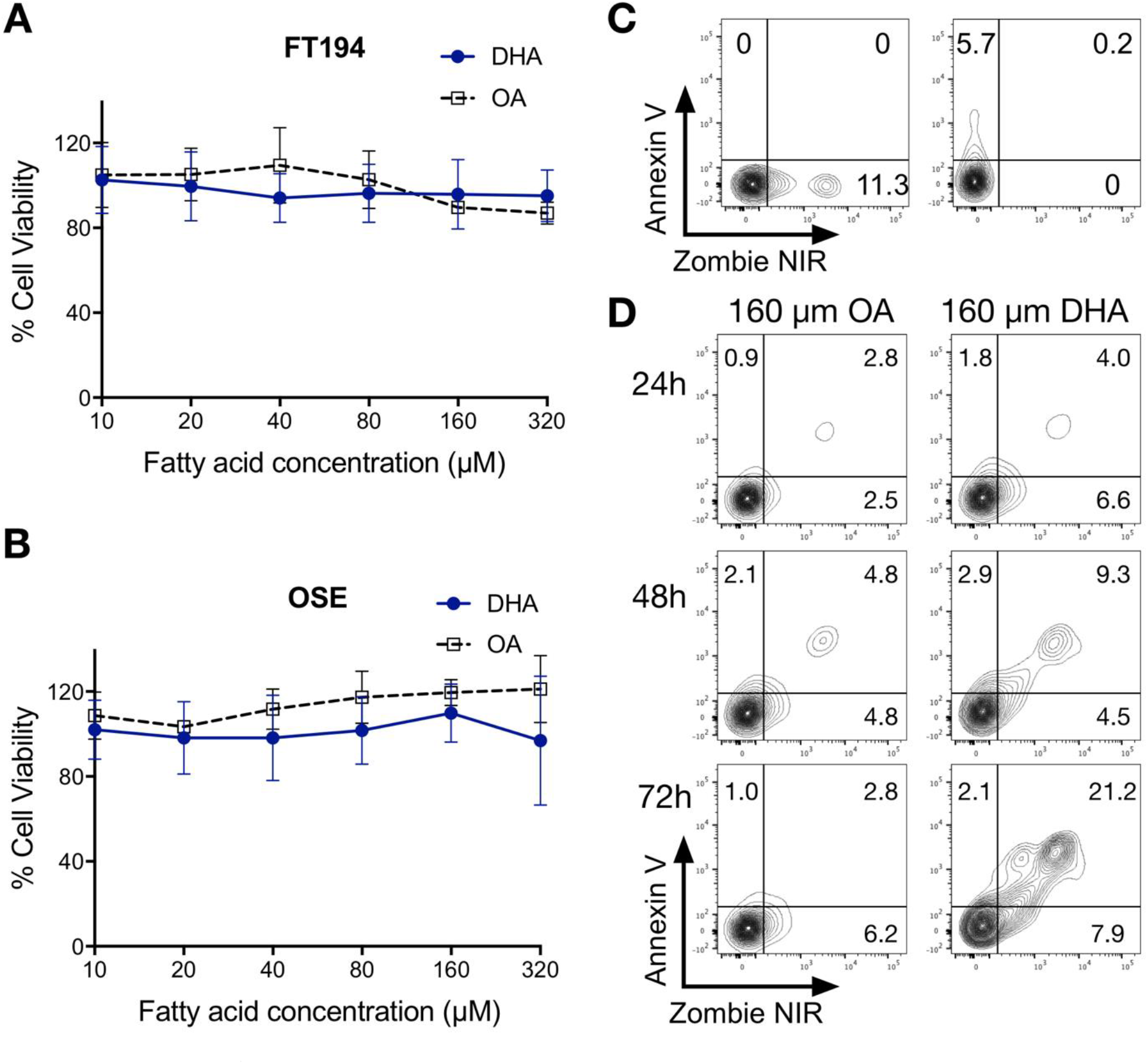
Effect of N-3 PUFAs DHA and OA on normal immortalised human fallopian tube FT194 cells and immortalized human ovarian surface epithelial cells IOSE-364 (OSE). (A and B) MTT assays showing cell viability of FT194 and OSE cells after treatment with OA and DHA at different concentrations for 72h. Treatment with 10-320 μM DHA or OA did not significantly reduce cell viability of either lines. Values represent the mean±SD. (C) FT194 cells were treated with 160 μM OA or DHA for 24, 48 and 72h, stained with Zombie Aqua and AnnexinV and acquired by flow cytometry to detect apoptosis. Fluorescence minus one (FMO) controls were used to set quadrant gates for contour plots of AnnexinV versus Zombie NIR. (D) DHA induced apoptosis in 21.1% FT194 cells after 72h treatment compared to 2.8% cells after treatment with OA.

**Supplementary Figure S5.**
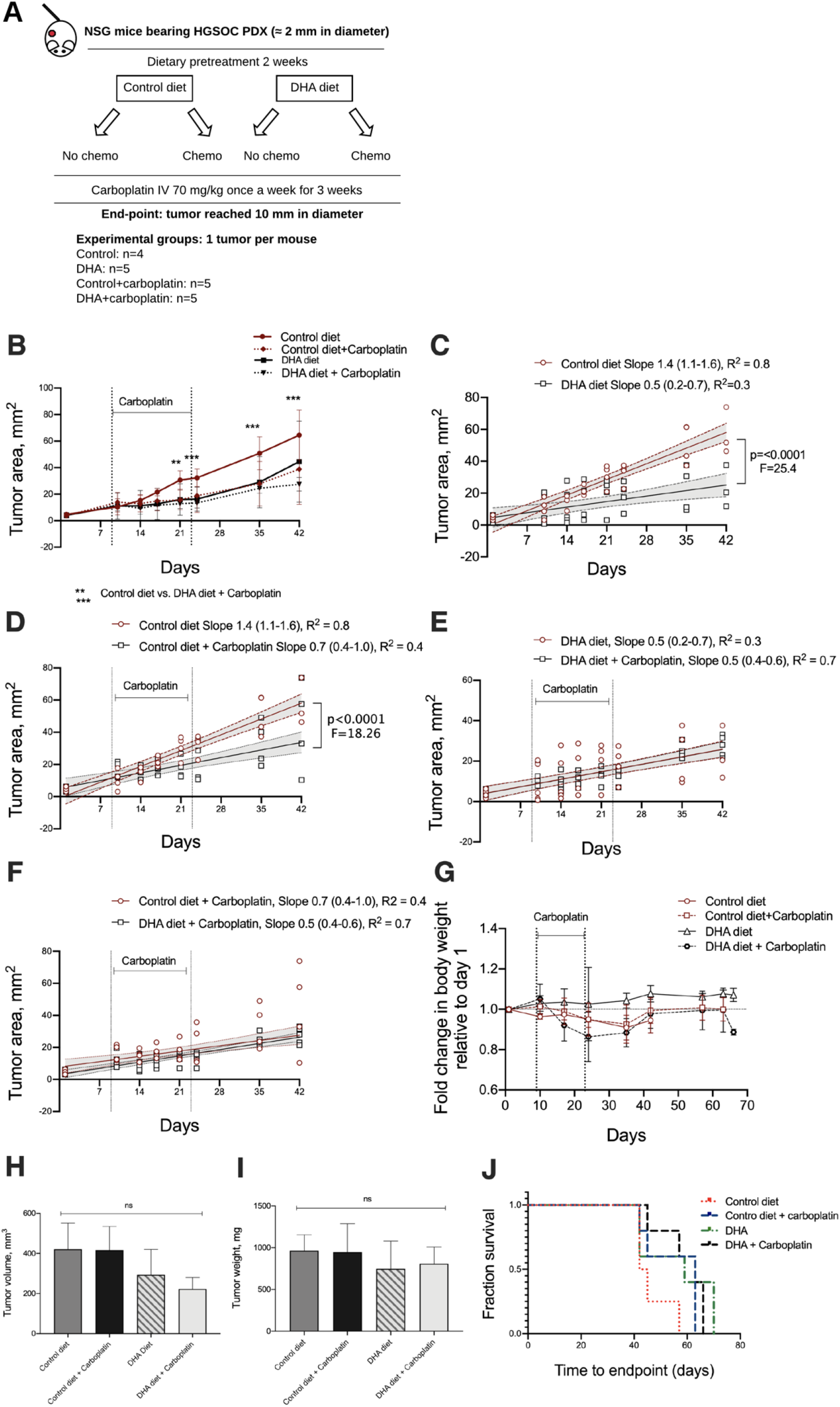
Effect of dietary DHA with or without intravenous (IV) carboplatin administration on HGSOC PDX-550 growth in NSG mice. (A) Experimental design: mice were implanted subcutaneously with HGSOC PDX-550 tumor sections approximately 2 mm in diameter. Two weeks prior to commencing chemotherapy, the mice were randomized into control or DHA diet groups and subsequently further randomized into carboplatin chemotherapy or vehicle control *via* intravenous (IV) administration. Carboplatin administration started when tumors reached approximately 10-20 mm^2^. Experimental groups are defined as control diet, control diet + carboplatin, DHA diet, and DHA diet + carboplatin. The experimental endpoint was defined as the point at which the tumor reached 10 mm in diameter (~60 mm^2^). (B) Average tumor area of HGSOC PDX-550 tumor bearing mice over time. Values represent the mean±SD. ** statistical difference between day 21 of control compared to DHA + carboplatin, *** statistical difference between day 24 of control compared to DHA + carboplatin. (C) Non-linear regression curve fit of straight line showing significant difference in slopes of tumor growth between control and DHA diet groups, (D) control diet and control diet + carboplatin groups. (E, F) Non-linear regression curve fit of straight line showing no difference in tumor growth between DHA and DHA + carboplatin group, control + carboplatin and DHA + carboplatin groups. Values in non-linear regression curve fit represent individual replicates with 95% CI. (G) Fold change in body weight of mice randomized into different groups relative to day one. There were no significant differences between groups. (H) Excised tumor volume and (I) tumor weight of PDX tumor bearing mice. (J) Kaplan-Meier curves comparing overall survival among groups.

**Supplementary Figure S6.**
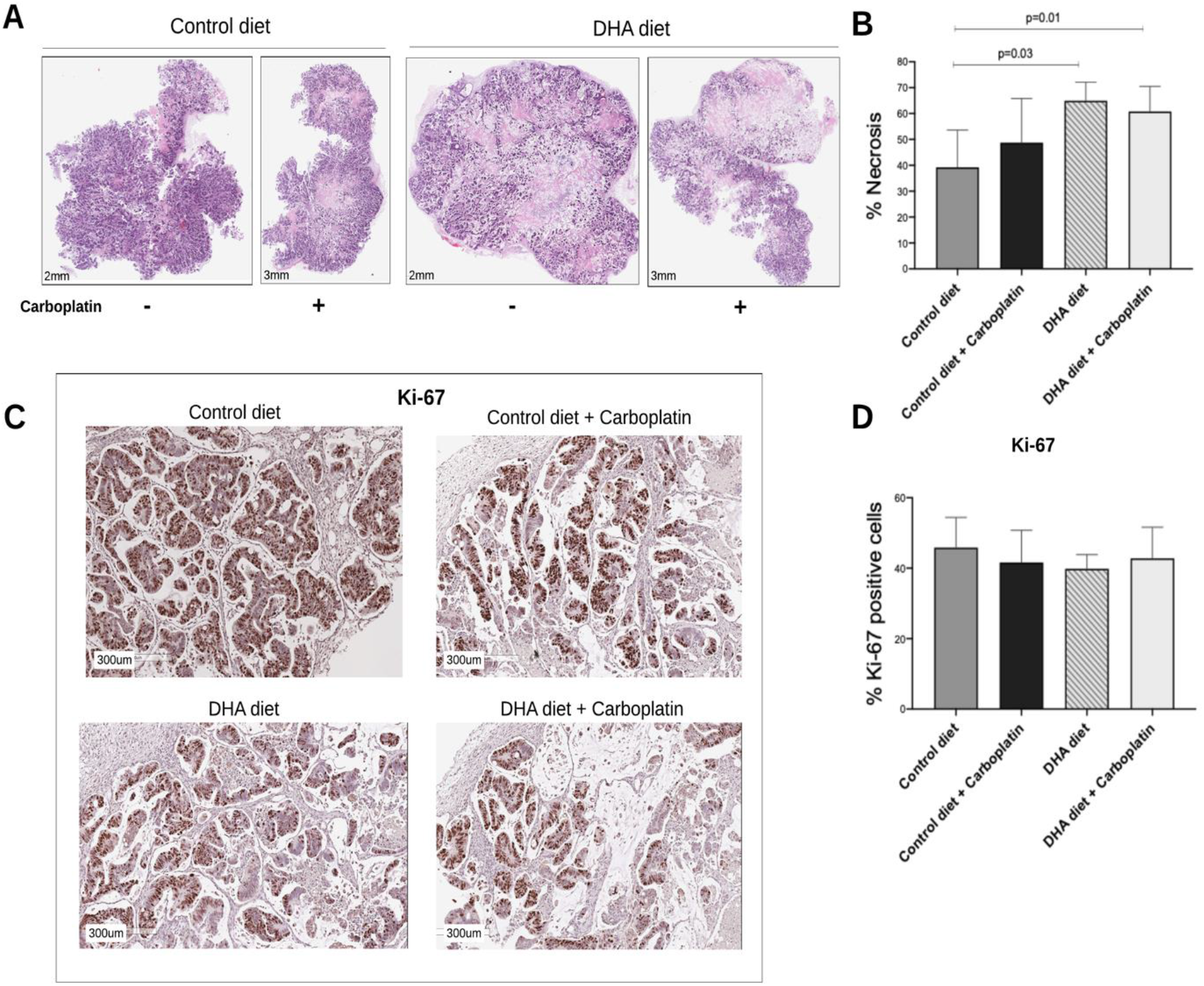
Effect of dietary DHA with or without intravenous (IV) carboplatin administration on HGSOC PDX-550 tumor necrosis and cell proliferation in NSG mice. (A) Representative H&E images of extracted tumors from NSG mice randomized into control or DHA groups with or without carboplatin administration. (B) Average area of necrosis in tumor extracts from NSG mice randomized into different groups. Tumors extracted from DHA and DHA + carboplatin randomized groups show significantly larger areas of necrosis compared to tumors from mice on the control diet (p<0.05). (C) Representative images of proliferation marker Ki-67 IHC staining of tumor sections. (D) Number of cells expressing Ki-67 in tumor sections from NSG mice randomized into different groups. Proliferation rates in tumors extracted from different groups do not differ. Values represent mean±SD.

